# Prediction of standard cell types and functional markers from flow cytometry gating definitions using machine learning

**DOI:** 10.1101/2021.08.25.457664

**Authors:** Raul Rodriguez-Esteban, José Duarte, Priscila C. Teixeira, Fabien Richard, Svetlana Koltsova, W. Venus So

## Abstract

**Background:** A key step in clinical flow cytometry data analysis is gating, which involves the identification of cell populations. The process of gating produces a set of reportable results, which are typically described by gating definitions. The non-standardized, non-interpreted nature of gating definitions represents a hurdle for data interpretation and data sharing across and within organizations. Interpreting and standardizing gating definitions for subsequent analysis of gating results requires a curation effort from experts. Machine learning approaches have the potential to help in this process by predicting expert annotations associated with gating definitions.

**Methods:** We created a gold-standard dataset by manually annotating thousands of gating definitions with cell type and functional marker annotations. We used this dataset to train and test a machine learning pipeline able to predict standard cell types and functional marker genes associated with gating definitions.

**Results:** The machine learning pipeline predicted annotations with high accuracy for both cell types and functional marker genes. Accuracy was lower for gating definitions from assays belonging to laboratories from which limited or no prior data was available in the training. Manual error review ensured that resulting predicted annotations could be reused subsequently as additional gold-standard training data.

**Conclusions:** Machine learning methods are able to consistently predict annotations associated with gating definitions from flow cytometry assays. However, a hybrid automatic and manual annotation workflow would be recommended to achieve optimal results.

## Introduction

Flow cytometry enables high content analysis of cell populations from heterogeneous samples through the identification of surface and intracellular antigen expression using fluorescent-labeled molecular probes (Chattopadhyay et al., 2008). It can provide insights in applications such as the identification of disease biomarkers, immune regulatory mechanisms and cellular signaling. As such, flow cytometry is an important tool in drug discovery and development in areas such as biomarker discovery, receptor occupancy and target engagement assays, and target-based and phenotypic screenings (Gedye et al., 2014; Edwards & Sklar, 2015; Moulard & Ozoux, 2016).

Recent years have seen tremendous development in multiplexing capabilities of flow cytometry instrumentation, namely with the development of full spectrum flow cytometry (Robinson, 2019; Nolan et al., 2013). This innovation has reached the clinical space and is implemented in high parameter flow cytometry assays in multi-center clinical trials. Such trials generate data from hundreds to thousands of samples across multiple flow cytometry assays that are capable of reporting on hundreds to thousands of different reportable results (“reportables”). Leveraging these new capabilities, for instance, pharmaceutical companies employ an evolving mix of flow cytometry assays during the drug project life cycle, stemming from a range of internally-developed assays and potentially multiple external laboratories.

The sharing of the biomarker data produced by these assays enables its reuse, reanalysis and reproducibility (Bhattacharya et al., 2018). However, effective sharing following best practices, such as those exemplified by the FAIR data sharing principles (Wilkinson et al., 2016), presents numerous challenges, such as those deriving from differences in sample management, instrumental setup and data analysis (Maecker et al., 2010; Final et al., 2016; Montante et al., 2019). Although much attention has been given to the harmonization and alignment of flow cytometry instruments in multi-center trials (White et al., 2015; Jamin et al., 2016; Larbi, 2017; Finak et al., 2016), there is still no guidance and/or tools for the standardization of flow cytometry data analysis and harmonization, such as the management of an assay’s multiple reportables. Moreover, data inconsistencies and errors can make cross-study analysis particularly difficult to execute. These factors, among others, increase the burden of deploying high parameter flow cytometry in the clinic.

The focus of this study is on the description of reportables, which are typically described using unstructured text strings, sometimes referred to as “gating definitions,” that comprise relevant markers and other information about the assay. Due to a lack of widespread standards, gating definitions can be written in multiple ways, which is recognized as an obstacle for data sharing (Overton et al., 2019). Integration of gating definitions from flow cytometry datasets is typically done through manual curation, with its associated perils, such as curation inconsistencies, drift and errors (Rodriguez-Esteban, 2015).

Overton et al. (2019) introduced an approach to check the validity and consistency of gating definitions for 4,388 gating definitions produced by a set of 28 academic centers. Their approach leveraged ontology mapping and, in particular, the Cell Ontology (CL) (Diehl et al., 2016) and the Protein Ontology (PRO) (Natale et al., 2011; Chen et al., 2020). It involved, among other steps, mapping gating definitions to marker gene names and intensity levels using a rule-based method. As stated by the authors, however, pure rule-based approaches have shortcomings in dealing with textual ambiguity. Owing to incomplete ontologies, ontology mapping can lead to false negatives due to unmatched relevant concepts. Additionally, rule-based methods can struggle to capture complex relations between elements of the text.

In this study, we explored a related problem to that reported in Overton et al. (2019). We studied the feasibility of predicting cell types and functional markers associated with gating definitions with the help of machine learning. Functional markers are markers that provide additional properties (e.g. proliferation, exhaustion status) but are not needed to define the cell types of interest in a particular assay. The cell types associated with each gating definition in an assay and the presence or absence of specific functional markers are of key interest for analyses concerning flow cytometry data because the annotation of gating definitions with standard concepts enables data integration and re-use. To tackle this problem, we applied a supervised machine learning (ML) approach. ML approaches for handling unstructured text have long been deployed in pharmaceutical research and development, which involves the mining of large amounts of textual information (Rodriguez-Esteban, 2016). In particular, mapping and classification of unstructured text is an area in which ML algorithms for text mining have already shown their utility (Rodriguez-Esteban et al., 2019). Thus, in this paper, we wanted to explore the feasibility of automatically identifying cell types and functional markers from gating definitions using an ML algorithm.

## Methods

The dataset for this study comprised 4,849 gating definitions from 36 assay panels belonging to 4 different laboratories. Despite differences between the assays, some gating definitions were identical. After deduplication, we had a total of 3,045 unique gating definitions available.

Most of the unique gating definitions (*n*=3,043) were interpreted by scientific experts, which annotated them with 117 unique cell types and 70 unique functional markers. An example of annotation of interpreted cell type and functional marker can be seen in Table 1. These annotations initially lacked some consistency. That is, the same cell type or functional marker was written in different ways by different experts. To increase consistency of annotation, annotated cell types were mapped to an internal Roche cell type terminology which integrates domain experts’ feedback and multiple public ontologies including Cell Ontology, BRENDA Tissue Ontology, SNOMED, NCI Thesaurus and MeSH; and is hosted by the Roche Terminology System (RTS), which is an internally-developed platform for the management and distribution of highly curated terminologies currently covering around 130,000 concept entries, and which is completely built on a semantic technology stack providing uniform resource identifiers (URIs) to support data FAIRification at scale across all Roche functions and sites.Very briefly, the creation of a cell type concept in RTS depends on experimental evidence showing that the said cell type has cellular functions different from other existing cell types. The mapping to the RTS terminologies involved manual expert curation and, additionally, rule-based automated quality control. For example, annotated marker gene names were harmonized to CD names where available. After the harmonization, we had annotations corresponding to 56 unique cell types and 62 unique functional markers, which became the target variables for the ML algorithm.

**Table 1.** Examples of gating definitions mapped to cell types and markers.

| Gating definition | Cell type | Functional marker |
| --- | --- | --- |
| CD3+CD4+CD25_APC MFI | Lymphocyte T, CD4-positive | CD25 |
| CD8+PD-1 Tcell MESFNaHepLDTCL | Lymphocyte T, CD8-positive | CD279 |
| Median CTLA4 in CD19 | Lymphocyte B | CD152 |
| %4-1BB in CD16 | Natural killer cell | CD137 |

Generalizability and reproducibility were emphasized in building the overall ML prediction workflow. Gating definitions were pre-processed by transforming them to lowercase, eliminating non-ASCII characters and most non-alphanumeric characters. Following Overton et al. (2019), a simple set of rules was applied to split (“tokenize”) gating definitions into units by identifying “separator” elements. These units ( “tokens”) often corresponded to individual gates (e.g. “CD3+CD4+CD25+” was split into the tokens CD3, CD4 and CD25). Marker intensity definitions (e.g. plus and minus signs next to individual gates, such as + in “CD3+”), where they existed, were extracted for each token.

The dataset was divided into training (80%) and test (20%) sets. Features for ML were based on all unique tokens produced by tokenization of the training dataset. These features were then matched to all gating definitions in the training and test sets to produce, respectively, the training and test feature values. Matches were not allowed when there were numerical boundaries around the match (e.g. the feature 45ra matched the gating definition “CD45RA+” but the feature cd4 did not). Marker intensity definitions were used to further refine feature values (e.g. a minus sign led to a feature value of −1).

Ontology matching was applied to gating definitions to reduce feature-set cardinality. For this purpose, the Protein Ontology (PRO) release 62.0 in OWL format, which included 331,920 terms, was downloaded from the Protein Ontology Consortium site (proconsortium.org).

An ML pipeline was chosen with the aid of the TPOT autoML (automated ML) library. AutoML algorithms can help the end-to-end selection of an optimal pipeline of preprocessors, feature constructors, feature selectors, ML models and hyperparameter optimization for solving an ML task (He et al., 2021). The TPOT autoML library for classification (Olson & Moore, 2016; v. 0.11.7) selects a model from a list that, in its default configuration, includes Gaussian naïve Bayes, Bernoulli naïve Bayes, multinomial naïve Bayes, decision tree, extra trees, random forest, gradient boosting, K-nearest neighbors, linear support vector machine, logistic regression, extreme gradient boosting, stochastic gradient descent and multi-layer perceptron. The ML pipeline was selected by running the TPOT autoML algorithm on the training set. The selected pipeline was primarily evaluated in the test set, which had not been seen by the autoML algorithm. Secondarily, it was evaluated by 10-fold cross-validation on the entire dataset.

All code and data created for this study are available at: https://github.com/raroes/prediction-flow-cytometry-gating-definitions

## Results

A total of 3,043 gating definitions were manually annotated by scientific experts and harmonized to 56 unique cell types and 62 unique functional markers (Table 1). Using this dataset, we developed a machine learning (ML)-based prediction workflow to predict the cell type and functional marker annotation associated with each gating definition, i.e. to solve a 56-class and a 62-class classification problem, respectively.

For cell type annotation prediction, the data was split into training and test datasets. Based on the gating definitions in the training dataset, a total of 281 features were created through data pre-processing steps (see Methods). Feature values were extracted for both training and test datasets to feed the ML pipeline. An ML pipeline was selected and optimized by the TPOT autoML (automated machine learning) algorithm. This pipeline was based on a stacking architecture composed of a random forest classifier and a logistic regression classifier. Using this pipeline, prediction accuracy on the test set was 97.2%. The median AUROC (area under the curve of the receiver operating characteristic) for each class was 0.999 and the average AUROC was 0.95 ± 0.12. Low performance for classes with few available gating definitions in the training lowered the average AUROC. The histogram distribution of AUROC values for all cell type classes is shown in Figure 2. The overall 10-fold cross-validation accuracy for the ML pipeline was 94.2%.

Due to the way manually-curated cell type annotations were written, these could be split into segments separated by commas. E.g., “Lymphocyte T, CD8-positive, regulatory” could be split into 3 segments: “Lymphocyte T,” “CD8-positive” and “regulatory” (Table 1). An error analysis showed that, out of 177 classification errors made by the ML algorithm, 124 (70.1% of all errors) corresponded to discrepancies with the third segment of the annotations. E.g., predicting “Lymphocyte T, CD4-positive” when the actual class was “Lymphocyte T, CD4-positive, *naive*.” Most predictions were, on the other hand, correct with respect to the first segment, which corresponded to the broader cell type (e.g., monocyte, neutrophil, T-cell), except in 34 cases (19.2% of all errors).

Data availability by source (i.e. external laboratory or internal assay provider) varied widely, as can be seen in Table 2. To test the ability of an ML pipeline trained on data from one source to successfully make predictions on test data from another source (i.e. transfer learning), we performed several experiments. First, we tested a pipeline on data from a single source after being trained on the rest of the sources. Results of this experiment can be seen in the first column of Table 3 and indicate that lack of same-source data in the training set had a strong negative impact on ML pipeline performance. The second experiment involved mixing different amounts of same-source and other-source data in the training set. As can be seen in Table 3, the addition of more data to the training set, whether same-source or other-source, generally improved prediction accuracy. It can also be seen that including even as little as 10% of same-source data significantly increased prediction accuracy in most cases.

**Table 2.** Number and percentage of curated gating definitions available by source (i.e. external laboratory or internal assay provider).

| Source | % | n |
| --- | --- | --- |
| Internal | 3.0 | 92 |
| Laboratory 1 | 83.8 | 2551 |
| Laboratory 2 | 6.0 | 184 |
| Laboratory 3 | 7.1 | 216 |

**Table 3.** Accuracy of the prediction pipeline for cell types when tested on single-source data and trained on different sets of sources. The first column of results corresponds to a pipeline tested on 100% of the data available from a single source (same-source) and trained on data from the rest of the sources (other-source). The second column of results corresponds to an algorithm trained on 10% of same-source data plus 0% or 100% of other-source data. The third column of results corresponds to an algorithm trained on 50% same-source data plus 0% or 100% of other-source data.

| Training data | % of existing same-source | 0% | 10% |  | 50% |  |
| --- | --- | --- | --- | --- | --- | --- |
|  | % of existing other-sources | 100% | 0% | 100% | 0% | 100% |
| **Test data** | **Internal** | 66.3 | 25.3 | 83.1 | 89.1 | 91.3 |
|  | **Laboratory 1** | 26.9 | 84.4 | 88.5 | 98.1 | 98.1 |
|  | **Laboratory 2** | 58.2 | 47.6 | 63.3 | 62.0 | 75.0 |
|  | **Laboratory 3** | 24.5 | 42.1 | 59.0 | 87.0 | 92.6 |

We performed an analogous analysis with the prediction of functional marker annotations. We manually labeled 3,038 gating definitions with functional marker annotations corresponding to 62 unique marker names (Table 1). This dataset was then used for creating, training and testing an ML pipeline as previously described. An ML pipeline based on logistic regression with L2 regularization was selected by the TPOT autoML algorithm. The accuracy of this ML pipeline was 98.5% on the test set. The median AUROC was 1.00 and the AUROC average was 0.87 ± 0.32 (Figure 2). The overall 10-fold cross-validation accuracy was 95.0%.

We then carried out the same experiments that we had performed with cell type annotation prediction in order to test the performance of the ML pipeline when trained on data from a mix of sources (Table 4). Similarly to the case with cell types, an increase in training data availability, whether same-source or other-source, led to greater accuracy (with one exception) and small amounts of same-source data improved performance considerably.

**Table 4.**
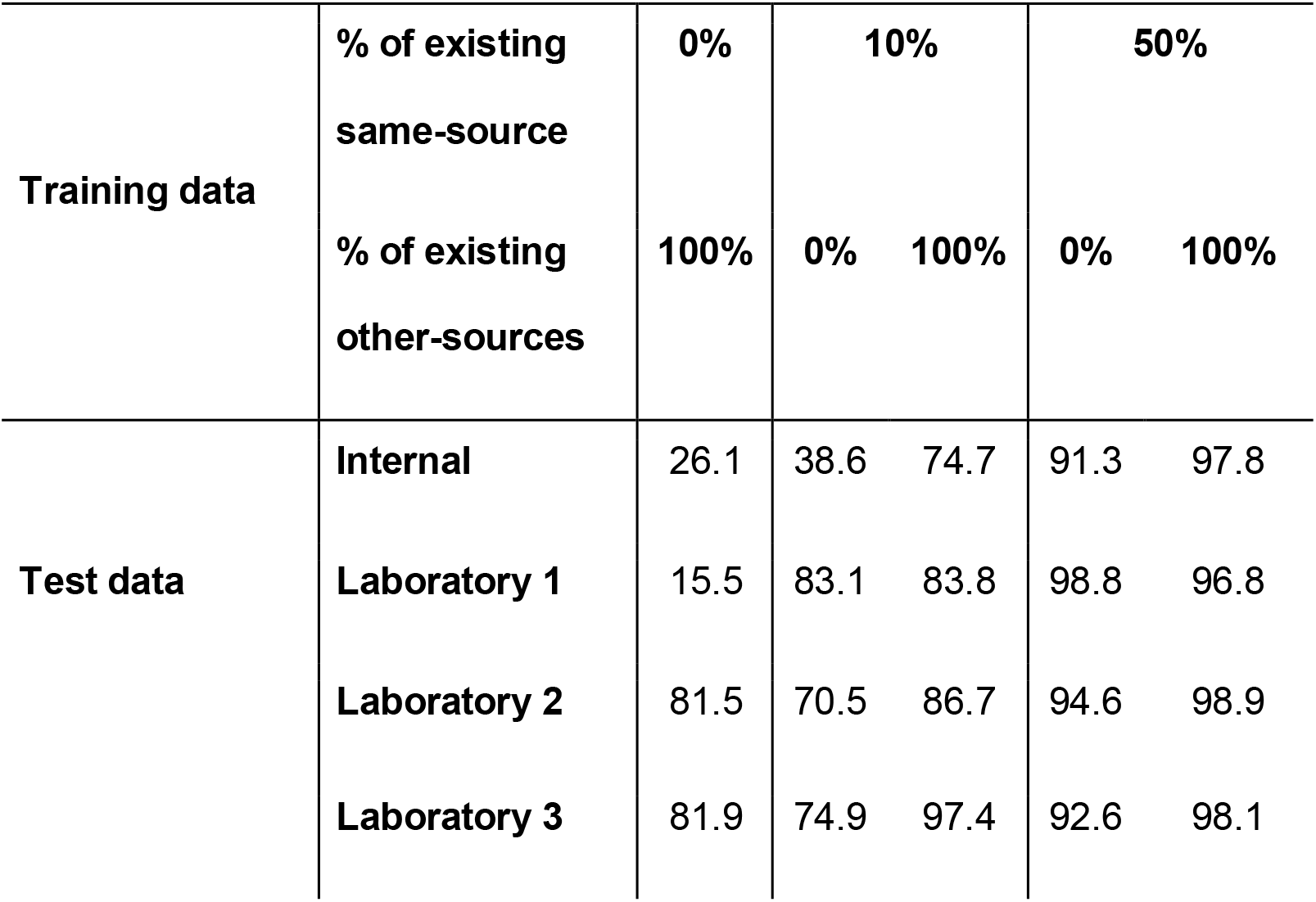
Accuracy of the prediction pipeline for markers when tested on single-source data and trained on different sources. The first column of results corresponds to an algorithm tested on 100% of the data available from a single source (same-source) and trained on data from the rest of the sources (other-source). The second column of results corresponds to an algorithm trained on 10% of same-source data plus 0% or 100% of other-source data. The third column of results corresponds to an algorithm trained on 50% same-source data plus 0% or 100% of other-source data.

Because gene names are highly ambiguous (Liu et al., 2006), we also explored following Overton et al. (2019) by using a gene name ontology (Protein Ontology, PRO) to identify features that could have been derived from different synonyms for the same gene name (see workflow in Figure 1). A total of 16 features identified in the training set that corresponded to synonyms from 8 genes were merged (e.g. the features corresponding to the gene name synonyms ICOS and CD278 were merged into one feature). This, however, did not improve performance for either cell type or marker classification.

**Figure 1.**
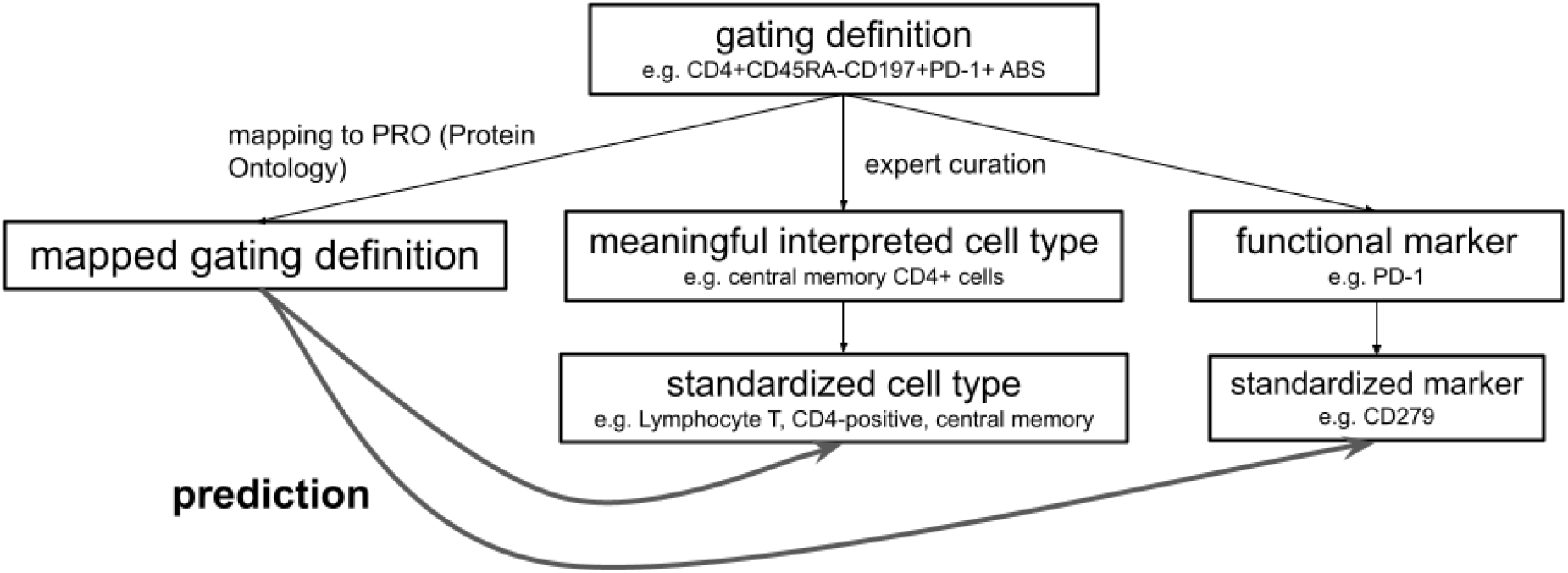
Prediction workflow for gating definitions mapped to ontology cell types.

**Figure 2.**
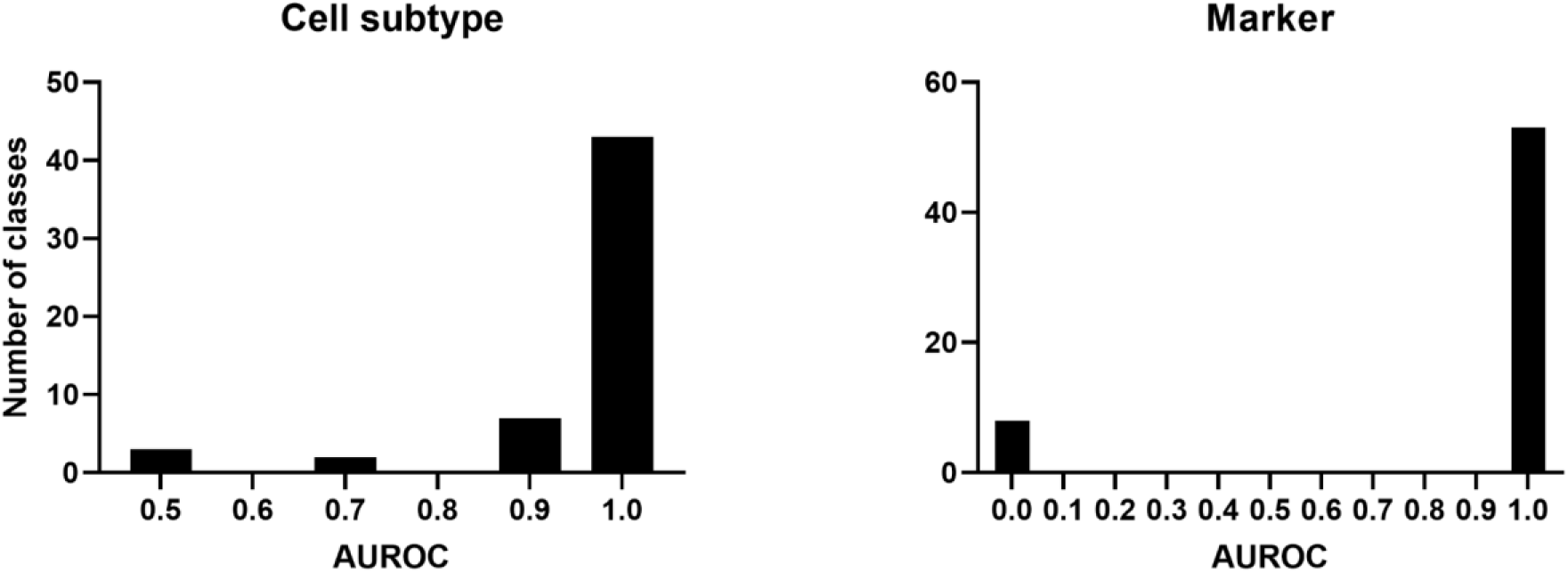
Histogram of AUROC values associated with the classification of each cell type and functional marker class. The abscissa is labeled with the upper bound of the histogram intervals.

## Discussion and conclusions

This study showed the feasibility of using ML algorithms for annotating flow cytometry gating definitions with standardized cell types and functional markers, thereby enabling the interpretation and integration of data provided by different assays deployed in multi-center studies. More accurate and efficient data integration increases the value of data and enhances its ability to generate clinical and biological insights. The ML algorithms chosen in this study can easily be re-trained as additional curated gating definitions are added to the training data.

Overall, the annotation of functional markers showed better performance than the annotation of cell types, which is possibly associated with differences in task complexity. While the training data that best helped predicting annotations belonged typically to assays developed by the same laboratory, we showed that the inclusion of gating definitions from assays belonging to other laboratories in the training data generally provided additional predictive power, thereby confirming the impact of transfer learning.

Cell type annotation errors were most frequent in fine-grained cell subtypes, as the main cell type was usually correctly predicted. As would be expected, annotating cell types and functional markers for which few examples were available in the training data was a challenge for the ML pipeline. This could be seen in the decreased AUROC for classes with low number of samples. Additionally, we observed that the annotation of gating definitions from assays belonging to laboratories for which there was no training data could lead to poor performance due to differences in the way gating definitions are written across laboratories. This can be addressed, as shown in our experiments, by curating a small set of representative gating definitions, so that the algorithm can learn to recognize the feature patterns that define data from a never-before-seen laboratory. Thus, while we have noted that “the more data the better,” manually-curated data can, nonetheless, be gathered strategically to increase its representativeness and improve the performance of the ML pipeline at low cost.

Additional challenges to our ML approach include those typical from operationalization of an ML algorithm (Mäkinen et al., 2021), such as tracking of ML model and dataset versions, as well as maintaining consistency in the quality of manual annotations. The ML algorithm itself can help in identifying consistency errors in manual annotations if an error analysis is performed on its predictions. Based on our practice, the output of the ML algorithm should be manually checked, which ensures high quality in the final output with minimal manual work. This output, in turn, can become additional high quality training data.

A clear advantage of a purely ML approach over a rule-based approach is that it does not depend on the currency, comprehensiveness or quality of the rules or ontologies used in the latter. However, the use of an ML approach does not preclude the inclusion of rules or ontologies. In fact, a mixed approach in which rules or ontologies were used to engineer features may improve performance. In our case, we tested the widely used ontology PRO to enhance our pipeline by normalizing features derived from synonyms corresponding to the same genes. This normalization did not, however, lead to an improvement in performance, perhaps due to the limited number of ambiguous gene synonyms identified.

## Acknowledgements

We would like to thank our managers Barbara Endler-Jobst and James Cai for their support. We would also like to thank our colleagues Iva Lelios, Fabian Junker and Enrique Gomez for their support with assay annotation, as well as our colleagues Nima Salimi and Yelena Budovskaya.

